# Muscle *FOXO*-specific overexpression and endurance exercise protects skeletal muscle and heart from defects caused by a high-fat diet in young *Drosophila*

**DOI:** 10.1101/2022.09.05.506594

**Authors:** Deng-tai Wen, Yi-ling Chen, Wen-qi Hou

## Abstract

Obesity appears to significantly reduce physical activity, but it remains unclear whether this is related to obesity-induced damage to skeletal muscle(SM) and heart muscle(HM). Endurance exercise(EE) reduces obesity-induced defects in SM and HM, but its molecular mechanism is poorly understood. The results showed that the structure and function of SM and HM were impaired by a high-fat diet(HFD) and muscle-FOXO-specific RNAi(MFSR), including reduced climbing speed and climbing endurance, reduced fractional shortening of the heart, damaged myofibrils, and reduced mitochondria in HM. Besides, a HFD and MFSR increased triglyceride level and MDA level, decreased the Sirt1 and FOXO protein level, and reduced CPT1, SOD, and CAT activity level, and they dow-regulated FOXO and bmm expression level in SM and HM. On the contrary, both muscle FOXO-specific overexpression(MFSO) and EE prevented abnormal changes of SM and HM in function, structure, or physiology caused by HFD and MFSR. Besides, EE also prevented defects of SM and HM induced by MFSR. Therefore, Current findings confirmed that MFSO and EE protected SM and heart from defects caused by a HFD via enhancing FOXO-realated antioxidant pathways and lipid catabolism. FOXO played a vital role in regulating HFD-induced defects in SM and HM, but FOXO was not a key regulatory gene of EE against damages in SM and HM. The mechanism was related to activity of Sirt1/FOXO/ SOD, CAT pathways and lipid catabolism in SM and HM.

## 1 Introduction

The childhood obesity epidemic has become a serious public health problem in many countries worldwide, and it is a major public health challenge of the 21st century[1]. In North America, approximately 30% of school age-children are overweight or obese, and 15% are obese. In European countries, these figures are 20% and 5%, respectively[2]. Besides, more than 20 million children and adolescents are overweight and obese in China[3]. Obesity in both adults and children is associated with numerous comorbidities, including muscle atrophy and cardiovascular diseases, and these obesity-induced diseases are closely related to the increase of lipotoxicity of vital organs in the body[4, 5]. Moreover, obesity can lead to adverse changes in the structure and function of skeletal muscles in children. For example, obesity results in increased adipose tissue and decreased skeletal muscle mass, and the accumulation of adipose tissue in skeletal muscle is closely related to insulin resistance and type 2 diabetes in children[6, 7]. Besides, the postural balance of obese children was impaired at different stable levels, and the calf muscles of obese children were less flexible[8]. As we all know, long-term HFD and lack of physical exercise are important causes of childhood obesity, and in children and adolescents, the more obese individuals are, the more difficult it is to participate in physical exercise[9, 10]. This phenomenon may be related to the physiological changes of skeletal muscle induced by obesity, but the underlying physiological mechanism is still poorly understood.

Forkhead O transcription factor (FOXO) and FOXO-related pathways may play a critical role in the regulation of lipid metabolism in heaert[11]. For instance, overexpression of *FOXO* gene in myocardial cells protects the heart from lipid accumulation and heart dysfunction induced by a HFD[12]. Besides, *brummer (bmm)* gene encodes a triglyceride lipase, which is involved in lipid metabolism, including regulation of lipid accumulation and triglyceride(TG) homeostasis[13]. Overexpression of *bmm* gene in heart prevents the heart from TG accumulation and heart defects induced by HFD[12]. Moreover, Sirtuin 1 (Sirt1) encodes an NAD^+^-dependent deacetylase that functions during euchromatic and heterochromatic gene silencing, and it plays a critical role in metabolic health by deacetylating many target proteins in numerous tissues, including heart, muscle, and adipose tissue[14–16]. Overexpression of *Sirt1* in heart prevents heart from TG accumulation, FOXO/SOD pathway inhabitation, and heart dysfunction induced by HFD or aging[17–19]. Malondialdehyde(MDA) is a product of lipid peroxidation, and this aldehyde is a highly toxic molecule and should be considered as a marker of lipotoxicity[20, 21]. Overexpression of *Sirt1* also decreases the MAD level by enhancing antioxidant capacity in aged heart[19]. Finally, physical exercise reduces lipid accumulation in skeletal muscle and the heart, and it increases their antioxidant capacity[22–26]. However, it remains unknown whether FOXO-specific expression in skeletal muscle and heart mediates exercise against defects induced by HFD in skeletal muscle and heart.

In this study, to determine whether FOXO-mediated antioxidant pathways are involved in regulating E against lipotoxicity induced by HFD in young muscle and heart, muscle or heart FOXO-specific overexpression (FSO) flies and FOXO-specific RNAi(FSR) flies were subjected to a HFD and PE intervention from day 3 and continued for 5 days, and then the function of muscle and heart, the lipid metabolic state, the antioxidant capacity, and the lipotoxicity level in both muscle and heart were measured by several methods.

## 2 Materials and methods

### 2.1 Fly stocks, diet and husbandry, and exercise training protocols

The *FOXO*-UAS-overexpression(*FOXO*-UAS-OE) flies (stock ID: 9575; FlyBase Genotype: y^1^ w^*^; P{UAS-foxo.P}2) [27], the *Mhc-gal4* (stock ID: 55133; FlyBase Genotype: w^*^; P[28]2/TM3, Sb^1^) flies, and *tin-gal4*(stock ID: 91538; FlyBase Genotype: y^1^ w^*^; P{tin-Gal4.B}2) were obtained from the Bloomington Stock Center. The *FOXO*-UAS-RNAi flies (stock ID: v106097; FlyBase Genotype: P {KK108590}VIE-260B) was obtained from the Vienna Drosophila Resource Center. “ P{KK108590} VIE-260B>w1118>MHC-gal4”, “P{UAS-foxo.P}2>w1118>MHC-gal4”, P{UAS-foxo.P}2>MHC-gal4”, and “P{KK108590} VIE-260B>MHC-gal4” were represented as “FOXO-RNAi-Control(FOXO-RNAi-C)”, “FOXO-OE-Control (FOXO-OE-C)”, “FOXO-over-expression (FOXO-OE)”, and “FOXO-RNAi” respectively. All UAS and GAL4 insertions were backcrossed into the w1118 line at least 10 times.

Normal food contained 1.6% soybean powder, 2.0% yeast, 6.7% corn meal, 0.7% agar, 4.8% sucrose, 4.8% maltose, and 0.3% propionic acid. Add 15% lard to normal food to make HFD[29, 30]. Flies were housed in a 25°C incubator with 50% humidity and a 12-h light/dark cycle. Fresh food was provided every other day.

The melting point of lard was 28-48°C. In our previous studies, we found that since coconut oil had a melting point of about 23 °C, and flies tended to stick to the melted coconut oil at 25°C, which leaded to an unnatural death in large numbers. However, the melting point of lard was 28-48°C. Flies could eat solid lard without sticking to it, which reduced unnecessary deaths.

When constructing the exercise device, the advantage of the flies’ natural negative geotaxis behavior was taken to induce upward walking[31]. Vials were rotated at the 60 rad/s. After vials each up-and-down turn, hold for 10 seconds for the flies to climb. Flies were exercised for 1.5 hours per day[32, 33]. All exercise groups flies started exercise from when they were 2-day old, and underwent a 6-day-long exercise program. All HFD flies started feeding high-diet food from when they were 2-day old, and underwent a 6-day-long high-fat diet program.

### 2.2 Climbing ability assay

The test vials for flies’ climbing speed and climbing failure were the same as the vials for exercise training, and the test vials were left eight centimeters length for the flies to climb[18]. With a light box behind the vials, once flies were shaken to the bottom of the vials, a timed digital camera snapped a picture after 3 seconds. The height the fly climbs was clearly shown on the photographs. Each vial contained about 20 flies and was subjected to 5 trials.

A cohort of flies was observed during continuous stimulation by the exercise device. Flies were placed on the exercise device in vials of 20 each and made to climb until fatigued. A vial of flies was considered “fatigued” when 5 or fewer flies are able to climb higher than 2 inches for four consecutive drops. Times of removal could be plotted as a “time-to-failure” plot [31].

### 2.3 Heart function assay

Flies were anesthetized with FlyNap for 2-3 min. The head, ventral thorax, and ventral abdominal cuticle were removed by special glass needles to expose the heart and abdomen. Dissections were done under oxygenated artificial hemolymph[34]. Artificial hemolymph was allowed to equilibrate with oxygenation for 15-20 min before filming. To get a random sampling of heart function, a single 30-s recording(AVI format) was made for each fly by using high-speed camera. The heart physiology of the flies was assessed using a AVS Video Editor analysis program that quantifies diastolic interval(DI), systolic interval(SI), heart period(HP), diastolic diameter(DD), systolic diameter(SD), and fractional shortening(FS)[35]. The sample size was 17 flies for each group.

### 2.4 MDA assay

20 flies muscle were converted to homogenates in a homogenizer filled with 1 ml PBS (pH 7.2-7.4). The homogenates were centrifuged at 4°C for 15 min with a speed of 2000 r/min. The supernatant was mixed with the reagents supplied in an MDA Assay Kit(MLBIO, Shanghai, China) and incubated at 95°C for 40 min. The absorbance of the supernatant was measured at 530 nm. All operations were according to the manufacturer’s instructions. All assays were repeated three times.

### 2.5 Transmission electron microscopy of skeletal muscle and myocardium

For electron microscopic analysis, skeletal muscle and myocardium were dissected in ice-cold fixative (2.5% glutaraldehyde in 0.1 M PIPES buffer at pH 7.4). After 10 hours of fixation at 4°C, samples were washed with 0.1M PIPES, post-fixed in 1% OsO4 (30 min), and stained in 2% uranyl acetate (1 hour). Samples were dehydrated in an ethanol series (50%, 70%, 100%) and embedded in epoxy. Ultrathin sections (50 nm) were cut and viewed on a Tecnai G2 Spirit Bio-TWIN electron microscope[36].

### 2.6 ELISA assay

The FOXO level, Sirt1 level, carnitine palmityl transferase-1(CPT1) level, SOD activity level, and CAT activity level were measured by ELISA assay(Insect FOXO, Sirt1, CPT1, SOD, and CAT ELISA Kits, MLBIO, Shanghai, China). 10 flies’ muscles and 80 hearts were homogenized in PBS (pH 7.2–7.4). Samples were rapidly frozen with liquid nitrogen and then maintained at 2°C - 8°C after melting. Homogenize the samples with grinders, and centrifugation was conducted for 20 min at 2000–3000 rpm. Then we removed the supernatant. Assay: take the blank well as zero, and read absorbance at 450 nm within 15 min of adding Stop Solution. All assays were repeated 3 times.

### 2.7 qRT-PCR

About 20 flies’ muscles and 80 hearts were homogenized in Trizol. 10 μg of the total RNA was purified by organic solvent extraction from the Trizol (TRIzol, Invitrogen). The purified RNA was treated with DNase I (RNase-free, Roche) and used to produce oligo dT-primed cDNAs (SuperScript II RT, Invitrogen), which were then used as templates for quantitative real-time PCR. The rp49 gene was used as an internal reference for normalizing the quantity of total RNAs. Real-time PCR was performed with SYBR green using an ABI7300 Real time PCR Instrument (Applied Biosystems), with 3 biological replicates. Expression of the various genes was determined by the comparative CT method (ABI Prism 7700 Sequence Detection System User Bulletin #2, Applied Biosystems). Primer sequences of *dFOXO* were as follows: F: 5’-AACAACAG CAGCATCAGCAG-3’; R:5’-CTGAA CCCGAGCATTCAGAT-3’. Primer sequences of *bmm* were as follows: F: 5′-ACTGCACATTTCGCTTACCC-3′, R: 5′-GAGAA TCCGGGTATGAAGCA-3′; Primer sequences of Rp49 were as follows: F: 5 –CTAAGCTG TCGCACAAATGG-3’; R: 5’-AACTT CTTGAATCC GGTGGG-3’.

### 2.8 Statistical analyses

Independent-sample *t* tests were used to assess differences between the each group flies. P-values for lifespan curves and climbing endurance curves were calculated by the log-rank test. Analyses were performed using the Statistical Package for the Social Sciences (SPSS) version 16.0 for Windows (SPSS Inc., Chicago, USA), with statistical significance set at P<0.05. Data are represented as means ± SEM.

## 3 Results

### 3.1 A 6-day HFD with 15% lard induces the defects of skeletal muscle(SM) in young *w*^*1118*^ wild *Drosophila*

It has been comformed that a 5-day HFD contributes to climbing performance decline, obesity phenotypes, and heart defects in flies, and this HFD is added to a normal diet with 30% coconut oil[17, 37]. Since coconut oil had a melting point of about 23 °C, and flies tended to stick to the melted coconut oil at 25°C, which leaded to an unnatural death in large numbers. To avoid large numbers of accidental fly deaths, we substituted 15% lard for 30% coconut oil for a HFD in this study. Increasing evidence conforms that the climbing speed and climbing endurance of flies can reflect its climbing performance[38].

The results showed that both a 2-day HFD, a 4-day HFD, and a 6-day HFD significantly decreased the climbing height(CH) in *w*^*1118*^ wild *Drosophila*(P<0.01, P<0.01, P<0.01) (Fig.1-A and B), and a 6-day HFD significantly decreased the time to fatigue(TTF) in *w*^*1118*^ wild *Drosophila* (P<0.01)(Fig.2-A). Besides, a 6-day HFD significantly increased the body weitht, body TG level, and muscle TG level(P<0.05, P<0.01, P<0.01) (Fig.2-B, E, G, and I), and it significantly reduced the body carnitine palmityl transferase-1(CPT1) in *w*^*1118*^ wild *Drosophila* (P<0.01) (Fig.2-F). These results suggeseted that a 6-day HFD with 15% lard impaired the climbing performance, contributed to obesity, and induced the reduction of key enzymes of lipid catabolism in young *w*^*1118*^ wild *Drosophila*.

**Fig.1.**
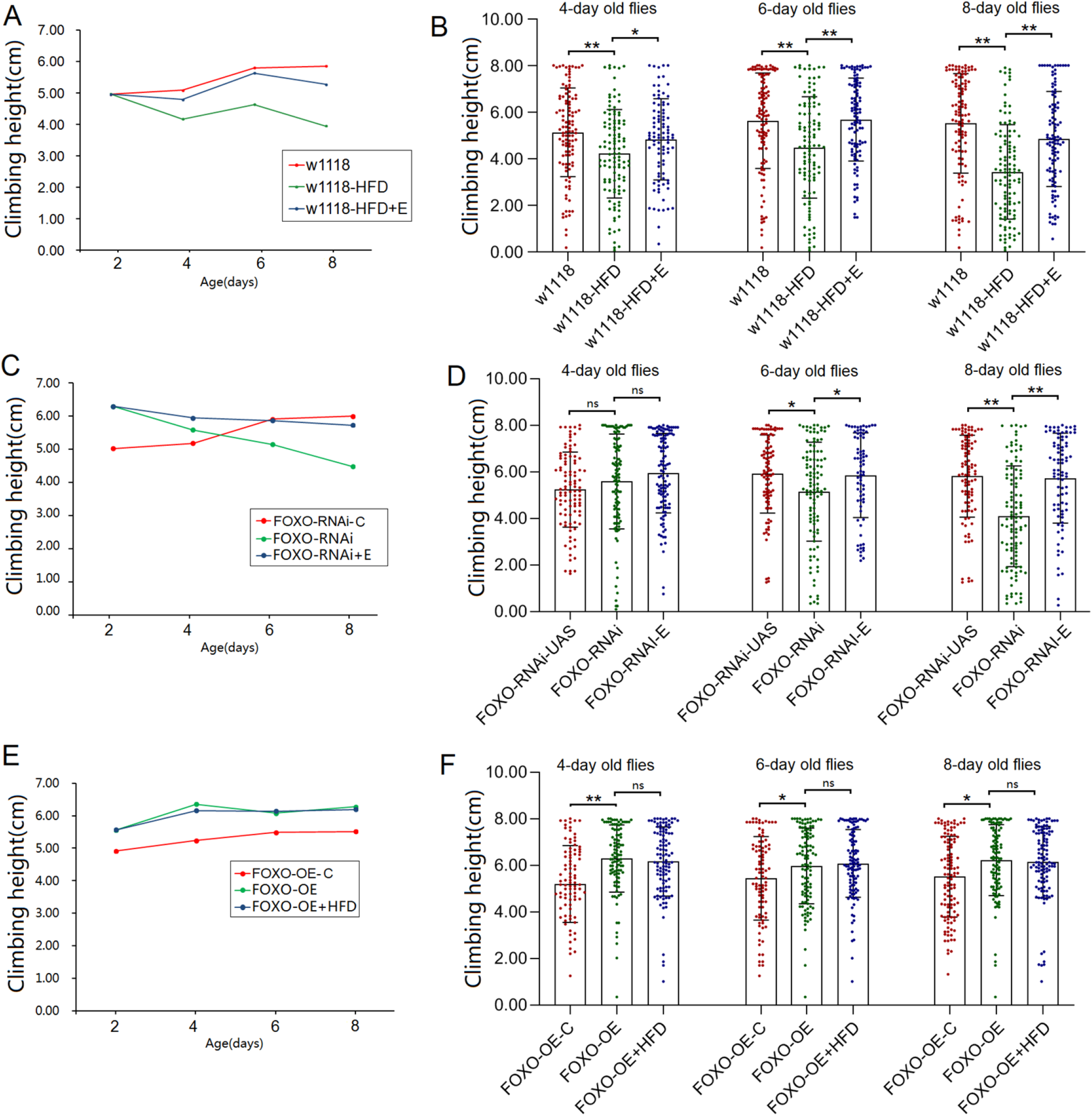
The climbing height in 3 seconds of flies. **(A)** Curve of climbing height(CH) in w^1118^ flies over 5 days. During those five days, the CH increased gradually, but a HFD caused the CH to decrease gradually, and exercise improved the CH reduction induced by a high-fat diet. (B) Comparison of CH in the same age group of w^1118^ flies. (C) Curve of CH in FOXO-RNAi flies over 5 days. During those five days, the CH increased gradually in FOXO-RNAi-UAS flies, but FSR caused the CH to decrease gradually, and exercise improved the climbing height reduction induced by FSR. (D) Comparison of CH in the same age group of FOXO-RNAi-UAS flies and FOXO-RNAi flies. (E) Curve of CH in FOXO-OE flies over 5 days. During those five days, the CH increased gradually in FOXO-OE-UAS flies and FOXO-OE flies, and HFD does not reduce CH in FOXO-OE flies. (F) Comparison of CH in the same age group of FOXO-OE-UAS flies and FOXO-OE flies. For CH measurement, the sample size was 100-120 flies for each group, and the 1-way analysis of variance (ANOVA) with least significant difference (LSD) tests was used to identify differences among the groups at the same age. Data are represented as means ± SEM. *P<0.05; **P <0.01.

**Fig.2.**
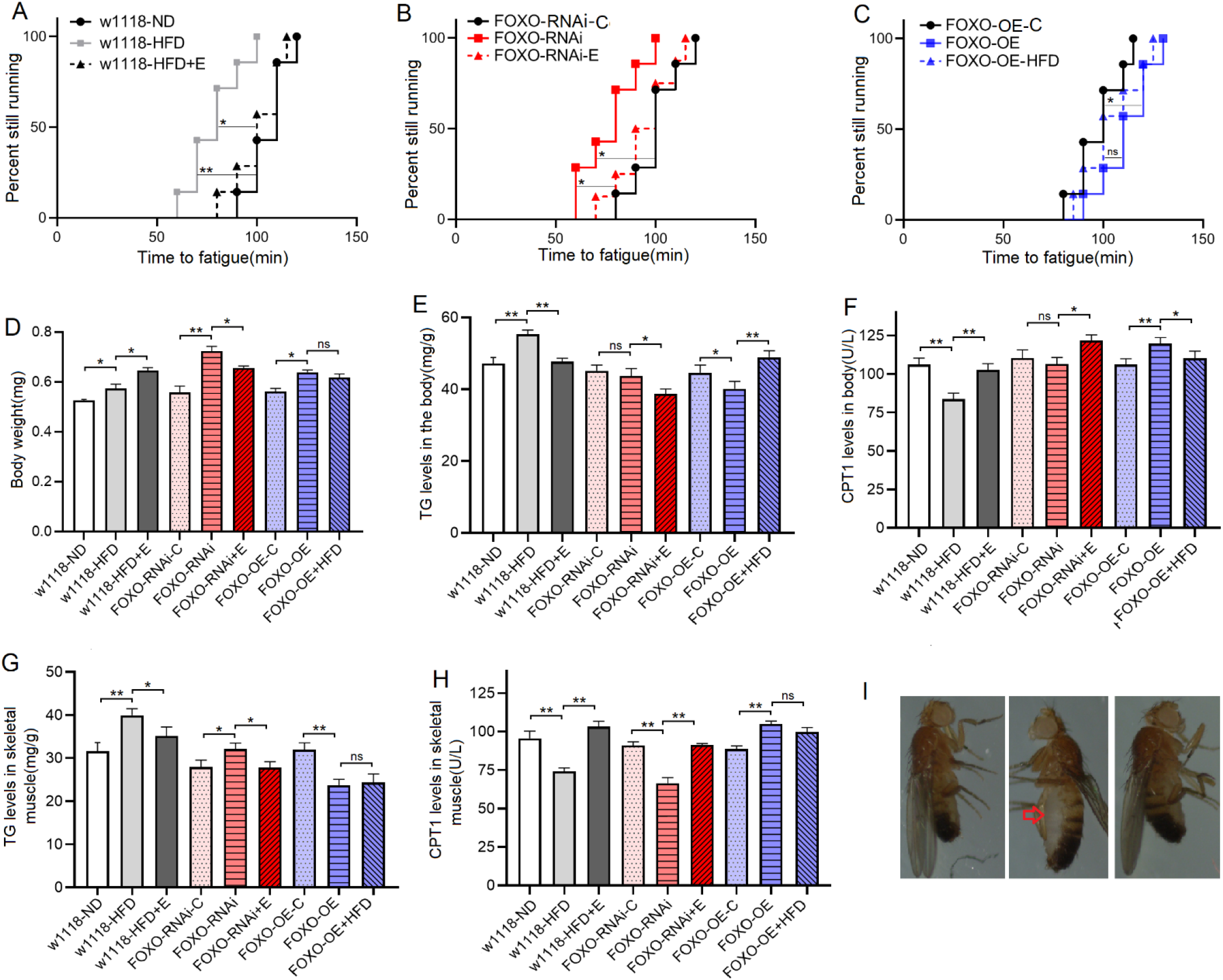
The climbing endurance and lipid metabolism of 8-day old flies. **(A)** Time to fatigue(TTF) in w^1118^ flies. (B) TTF in FOXO-RNAi flies. (C) TTF in FOXO-OE flies. (D) The body weight. (E) The body TG level. (F) The body CPT1 protein level. (F) The muscle TG level. (F) The muscle CPT1 protein level. (F) The images of w1118 flies. Red clipping indicated obvious lipid accumulation in abdomen induced by HFD. For climbing endurance, the sample size was 200-220 flies for each group, and P-values for climbing endurance curves were calculated by the log-rank test. For TG and CPT1 measurement, the sample size was about 20 flies’ skeletal muscle for each group, and the 1-way analysis of variance (ANOVA) with least significant difference (LSD) tests was used to identify differences among the groups with similar genetic backgrounds. Measurements were taken 3 times. Data are represented as means ± SEM. *P<0.05; **P <0.01.

To further investigate the changes in physiology induced by this HFD in SM, SM-related genes and proteins were detected by high-throughput sequencing, RT-PCR, and ELISA in young *w*^*1118*^ wild *Drosophila*[12, 37, 39, 40]. The high-throughput sequencing and RT-PCR results showed that the skeletal muscle *FOXO* expression was significantly dow-regulated by a 6-day HFD(P<0.01) (Fig.3-A to F). Besides, the results showed that the skeletal muscle CPT1 protein activity level and bmm expression level were significantly dow-regulated by a 6-day HFD(P<0.01, P<0.01) (Fig.2-H, Fig.3-F). Moreover, a 6-day HFD significantly reduced the skeletal muscle FOXO protein level, Sirt1 protein level, SOD activity level, and CAT activity level(P<0.01, P<0.01, P<0.01, P<0.05) (Fig.4-A to E), but it increased significantly skeletal muscle MDA level in young *w*^*1118*^ wild *Drosophila*(P<0.01) (Fig.4-F). In addition, the transmission electron microscope images displayed that a HFD induced myofibrillary damage (Fig.5).

**Fig.3.**
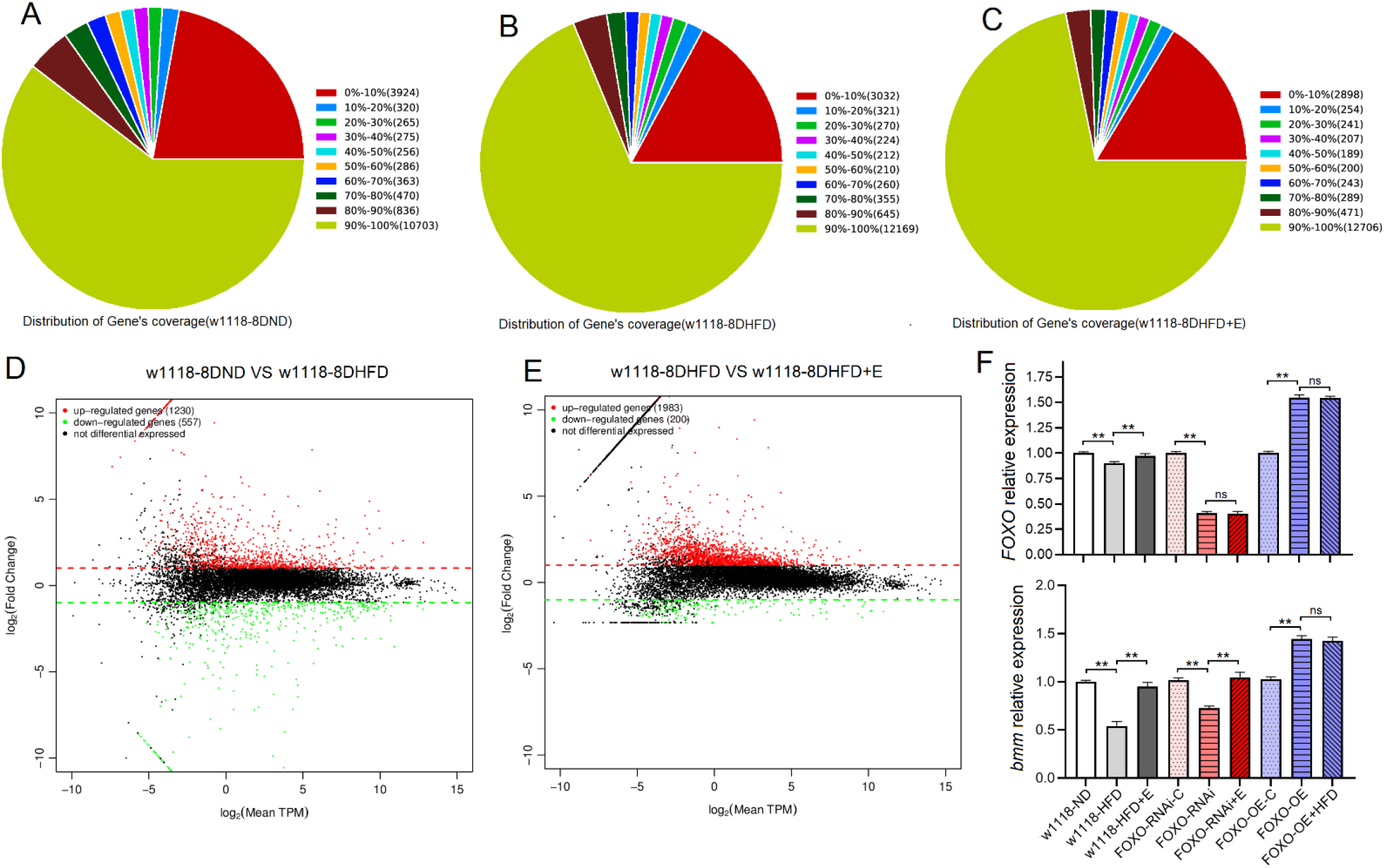
High-throughput sequencing and RT-PCR results of skeletal muscle of Drosophila w^1118^. **(A)** Distribution of gene’s coverage in w^1118^ flies. (B) Distribution of gene’s coverage in w^1118^-HFD flies. (C) Distribution of gene’s coverage in w^1118^-HFD+E flies. (D) Changes in gene expression in skeletal muscle w1118 compared with w1118-HFD. (E) Changes in gene expression in skeletal muscle w1118-HFD compared with w1118-HFD+E. (F)The expression of *FOXO* gene and *bmm* gene in skeletal muscle. High-throughput sequencing results showed that HFD significantly down-regulated the expression of *FOXO* gene and *bmm* gene, and exercise significantly upregulated *FOXO* gene and *bmm* gene. qRT-PCR results further confirmed the results of high throughput sequencing. For High-throughput sequencing and RT-PCR measurement, the sample size was about 30 flies’ skeletal muscle for each group, and the 1-way analysis of variance (ANOVA) with least significant difference (LSD) tests was used to identify differences among the groups with similar genetic backgrounds. Measurements were taken 3 times. Data are represented as means ± SEM. *P<0.05; **P <0.01.

**Fig.4.**
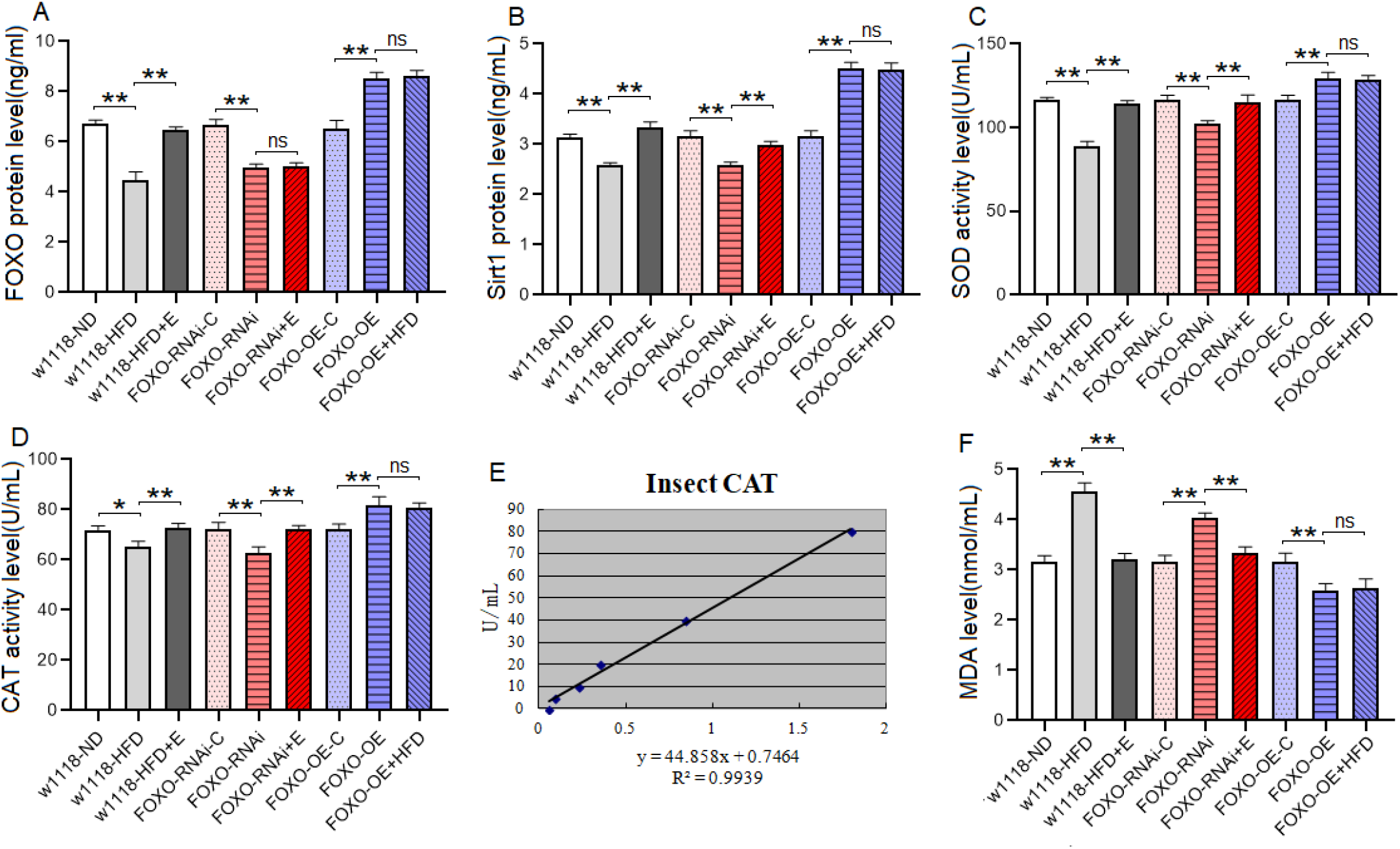
The Sirt1/FOXO pathways activity and lipotoxicity in skeletal muscle. **(A)** FOXO protein level. (B) Sirt1 protein level. (C) SOD protein activity level. (D) CAT protein activity level. (E) Standard curve of CAT test. (F) MDA level. For protein and MDA measurement, the sample size was about 20 flies’ skeletal muscle for each group, and the 1-way analysis of variance (ANOVA) with least significant difference (LSD) tests was used to identify differences among the groups with similar genetic backgrounds. Measurements were taken 3 times. Data are represented as means ± SEM. *P<0.05; **P <0.01.

**Fig.5.**
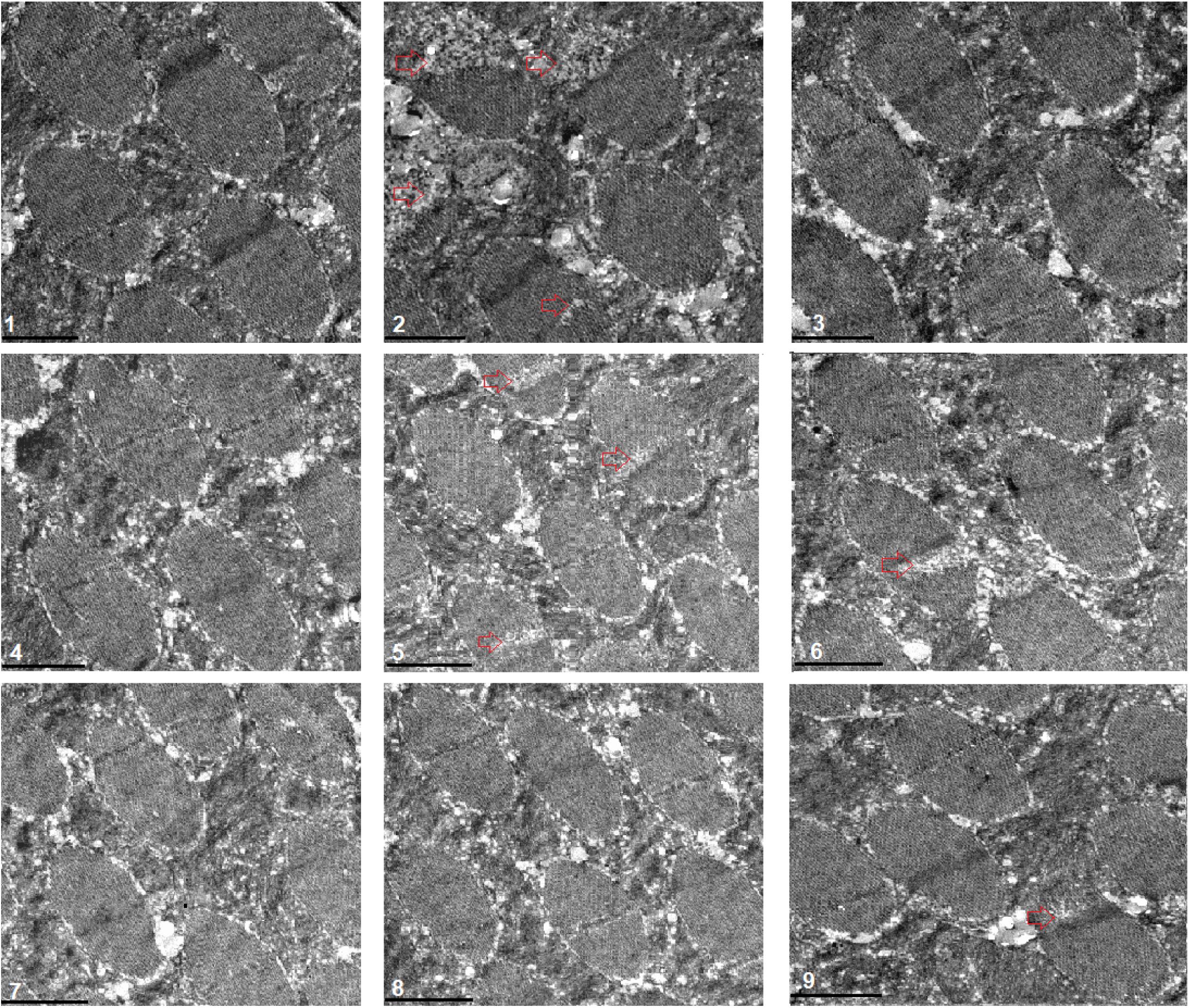
Transmission electron microscopy of muscle. 1: w^1118^ flies; 2: w^1118^-HFD flies; 3: w^1118^-HFD+E flies; 4: FOXO-RNAi-UAS flies; 5: FOXO-RNAi flies; 6: FOXO-RNAi+E flies; 7: FOXO-OE-UAS flies; 8: FOXO-OE flies; 9: FOXO-OE+HFD flies. Scale: the black line represents 2 microns. The transmission electron microscope images displayed that HFD and FSR induced myofibrillary damage, but PE could improve the myofibrillary damage induced by HFD and FSR. FSO protected myofibrillary from damage induced by HFD.

These results suggeseted that a 6-day HFD with 15% lard induced abnormal changes in lipid metabolism and antioxidant capacity, including the inhibition of CPT1 activity and bmm expression and dow-regulation of Sirt1/FOXO/SOD, CAT pathways, thus increasing the lipid accumulation in SM.

*Drosophila* has emerged as an important model to study the effects of HFD on metabolism, heart function, behavior, and ageing[39, 41–43]. For example, a 5-day HFD increases TG levels and disrupts insulin/glucose homeostasis in flies, which is similar to mammalian responses[44, 45], and this is consistent with our findings. It has been reported that the cardiac FOXO *gene* plays a key role in the regulation of lipid metabolism in the adult *Drosophila* heart[12], but few have reported the role of *FOXO* gene in the regulation of lipid metabolism in SM. The above experimental results indicate that FOXO gene and its related pathways may also be involved in the regulation of lipid metabolism in SM, but the role of SM-FOXO gene in regulating skeletal muscle lipid metabolism still needs to be further confirmed.

### 3.2 Skeletal muscle(SM) FOXO gene plays a key role in regulating skeletal muscle function and lipid metabolism

To comfirmed that the role of FOXO gene in regulating ipid metabolism in SM, muscle *FOXO-*specific overexpreesion(MFSO) and RNAi(MFSR) were built by UAS/MHC-Gal4 system. The results showed that MFSR significantly decreased the CH at age of 6-day old and 8-day old *Drosophila*(P<0.05, P<0.01) (Fig.1-C and D), and it also significantly decreased TTF at the age of 8-day old *Drosophila*(P<0.05) (Fig.2-B). Besides, the results showed that MFSR significantly increased body weitht(P<0.01) (Fig.2-D), but it did not induce a significant increase in whole-body TG(P>0.05) (Fig.2-E), and it also did not significantly decreased body CPT1 activity(P>0.05) (Fig.2-F). Moreover, the results showed MFSR significantly increased the skeletal mucle TG level(P<0.05) (Fig.2-G) and decreased CPT1 activity level(P<0.01) (Fig.2-H), FOXO gene expression and protein level(P<0.01, P<0.01) (Fig.3-F, Fig.4-A), bmm gene expression level(P<0.01) (Fig.3-F), Sirt1 protein level(P<0.01) (Fig.4-B), and SOD activity level and CAT activity level(P<0.01, P<0.01) (Fig.4-C to E), and MFSR significantly increased the skeletal mucle MDA level(P<0.01) (Fig.4-F). In addition, the transmission electron microscope images displayed that a MFSR impaired myofibrillary of skeletal muscle (Fig.5).

These results suggested that although FOXO knockdown in SM did not induce dysregulation of systemic lipid metabolism, muscle FOXO-specific knockdown in SM can lead to decrease in its function and abnormal lipid metabolism, including impaired climbing ability, increased lipid accumulation, attenuated lipid catabolism and antioxidant, which is similar to the effect of a HFD on SM. These results also suggested that FOXO gene played an important role in regulating lipid metabolism and antioxidant in SM, but whether this role is critical remains to be confirmed.

To further confirm whether FOXO gene played a key role in the regulation of lipid metabolism in SM, SM-FOXO gene was specifically overexpressed and the transgenic flies were subjected to a HFD intervention. The results showed that MFSO significantly increased the CH of 4-day-old, 6-day-old, and 8-day-old flies(P<0.01, P<0.05, P<0.05)(Fig.1-E and F), and it also significantly increased TTF of 8-day-old *Drosophila*(P<0.05) (Fig.2-C). Besides, the results showed that MFSO significantly increased body weight(P<0.05) (Fig.2-D) and whole-body TG level(P>0.05) (Fig.2-E), and it also significantly increased body CPT1 activity(P<0.01) (Fig.2-F). Moreover, the results showed MFSO significantly decreased the skeletal mucle TG level(P<0.05) (Fig.2-G) and increased CPT1 activity level(P<0.01) (Fig.2-H), FOXO gene expression and protein level(P<0.01, P<0.01) (Fig.3-F, Fig.4-A), bmm gene expression level(P<0.01) (Fig.3-F), Sirt1 protein level(P<0.01) (Fig.4-B), and SOD activity level and CAT activity level(P<0.01, P<0.01) (Fig.4-C to E), and MFSO significantly decreased the skeletal mucle MDA level(P<0.01) (Fig.4-F).

However, in MFSO transgenic flies, a 6-day HFD could not significantly change CH (P>0.05)(Fig.1-E and F), TTF(P>0.05) (Fig.2-C), skeletal mucle TG level(P>0.05) (Fig.2-G), CPT1 activity level(P>0.05) (Fig.2-H), FOXO gene expression and protein level(P>0.05, P>0.05) (Fig.3-F, Fig.4-A), bmm gene expression level(P>0.05) (Fig.3-F), Sirt1 protein level(P>0.05) (Fig.4-B), and SOD activity level and CAT activity level(P>0.05, P>0.05) (Fig.4-C to E), and skeletal mucle MDA level(P>0.05) (Fig.4-F). In addition, the transmission electron microscope images displayed that a MFSO protected myofibrillary of skeletal muscle from damage caused by a HFD (Fig.5).

These results suggested that FOXO-specific overexpression in skeletal muscle can enhance skeletal muscle function and lipid metabolism, including enhanced climbing ability and lipid catabolism, decreased lipid accumulation, and activated Sirt1/FOXO/SOD, CAT pathways related to antioxidant. In addition, the results showed that in MFSO transgenic flies, a 6-day HFD could not significantly reduce climbing ability, attenuate lipid catabolism, and inhibit Sirt1/FOXO/SOD, CAT pathways in SM. Therefore, the present results confirmed that in skeletal muscle, FOXO gene played a key role in preventing damages induced by a HFD, and its mechanism may be related to regulation of lipid catabolism and Sirt1/FOXO/SOD, CAT pathways related to antioxidant.

### 3.3 Endurance exercise(EE) prevented SM from damages induced by a HFD or MFSR in young *Drosophila*

Exercise training improves the climbing performance in adult flies, and it delays age-related climbing performance decline in aging flies[46–49]. Increasing evidence confirmes that a 5-day EE reduces lipid accumulation in SM and HM, and it increases their antioxidant capacity in flies[22–26], but the molecular pathways through which EE regulates skeletal muscle lipid metabolism are poorly understood.

In this study, the results suggested that a 2-day EE, 4-day EE, and 6-day EE significantly increased CH of in w^1118^-HFD flies (P<0.05, P<0.01, P<0.01)(Fig.1-A and B), and it also significantly increased TTF of 8-day-old HFD *Drosophila*(P<0.05) (Fig.2-A). Besides, the results showed that in *w*^*1118*^-HFD flies, a 6-day EE significantly increased body weight (P<0.05) (Fig.2-D), it significantly decresed whole-body TG level(P<0.01) (Fig.2-E), and it also significantly increased body CPT1 activity(P<0.01) (Fig.2-F). Moreover, the results showed that in *w*^*1118*^-HFD flies, a 6-day EE significantly decreased the skeletal mucle TG level(P<0.05) (Fig.2-G), and it significantly increased CPT1 activity level(P<0.01) (Fig.2-H), FOXO gene expression and protein level(P<0.01, P<0.01) (Fig.3-F, Fig.4-A), bmm gene expression level(P<0.01) (Fig.3-F), Sirt1 protein level(P<0.01) (Fig.4-B), and SOD activity level and CAT activity level(P<0.01, P<0.01) (Fig.4-C to E), and a 6-day EE significantly decreased the skeletal mucle MDA level(P<0.01) (Fig.4-F). In addition, the transmission electron microscope images displayed that EE protected myofibrillary of SM from damage caused by a HFD (Fig.5). These results suggested that EE could prevent SM-defects induced by a HFD through activating muscle lipid catabolism and Sirt1/FOXO/SOD, CAT pathways, but the role of SM-FOXO gene in this process remains unclear.

To confirm whether the FOXO gene plays a critical role in EE resistance to damages caused by lipid accumulation in SM, SM-FOXO-RNAi flies were subjected to EE intervention. The results showed that in SM-FOXO-RNAi flies, a 4-day EE and a 6-day EE significantly increased CH(P<0.05, P<0.01)(Fig.1-C and D), and it also significantly increased TTF (P<0.05) (Fig.2-B). Besides, the results showed that in SM-FOXO-RNAi flies, a 6-day EE significantly decreased body weight (P<0.05) (Fig.2-D), it significantly decresed whole-body TG level(P<0.01) (Fig.2-E), and it significantly increased body CPT1 activity(P<0.01) (Fig.2-F). Moreover, the results showed that in skeletal muscle FOXO-RNAi flies, a 6-day EE significantly decreased the skeletal mucle TG level(P<0.05) (Fig.2-G), and it significantly increased CPT1 activity level(P<0.01) (Fig.2-H), bmm gene expression level(P<0.01) (Fig.3-F), Sirt1 protein level(P<0.01) (Fig.4-B), and SOD activity level and CAT activity level(P<0.01, P<0.01) (Fig.4-C to E), and a 6-day EE significantly decreased the skeletal mucle MDA level(P<0.01) (Fig.4-F). However, in SM-FOXO-RNAi flies, a 6-day EE did not significantly change FOXO gene expression and protein level(P>0.05, P>0.05) (Fig.3-F, Fig.4-A). In addition, the transmission electron microscope images displayed that a MFSO protected myofibrillary of skeletal muscle from damage caused by skeletal muscle FOXO-RNAi (Fig.5).

These results suggested that EE could prevent defects induced by SM-FOXO-RNAi through activating muscle lipid catabolism and Sirt1/FOXO/SOD, CAT pathways, and FOXO gene was not a key gene for EE against damages induced by lipid accumulation in SM.

### 3.4 EE and MFSO prevented heart dysfunction induced by a HFD(with 15% lard) in young male *Drosophila*

It has been confirmed that cardiac *FOXO* gene play a vital role in regulating cardiac lipid metabolism and heart function, cardiac *FOXO* overexpression protects heart function from a HFD(with 30% coconut oil)-induced damage in adult drosophila flies, and cardiac *FOXO* RNAi can induce cardiac dysfunction, and this is similar to HFD(with 30% coconut oil)-induced cardiac dysfunction[12, 50],[12, 51, 52]. However, it remains unclear whether HFD(with 15% lard) could also cause cardiac dysfunction, and the role of FOXO gene in EE against lipid metabolism abnormalities and cardiac dysfunction in heart still remains unclear.

The results showed that at 8 days of age, the diastolic interval(DI) and heart period(HP) of w1118-HFD flies was significantly shorter than that of w1118 flies(P<0.05), and the DI and HP of w1118-HFD+E flies was significantly longer than that of w1118-HFD flies(P<0.05). Besides, the DI and HP of FOXO-RNAi flies was significantly shorter than that of FOXO-RNAi-C flies(P<0.05), and the DI and HP of FOXO-RNAi+E flies was significantly longer than that of FOXO-RNAi flies(P<0.05). Moreover, the DI and HP of FOXO-OE flies was significantly longer than that of FOXO-OE-C flies(P<0.05), but there was no significant difference in DI and HP between FOXO-OE flies and FOXO-OE+HFD flies(P>0.05). However, the effect of EE, HFD, FSO, and FSR on systolic interval was not significant (P>0.05). (Fig.6-A to C)

**Fig.6.**
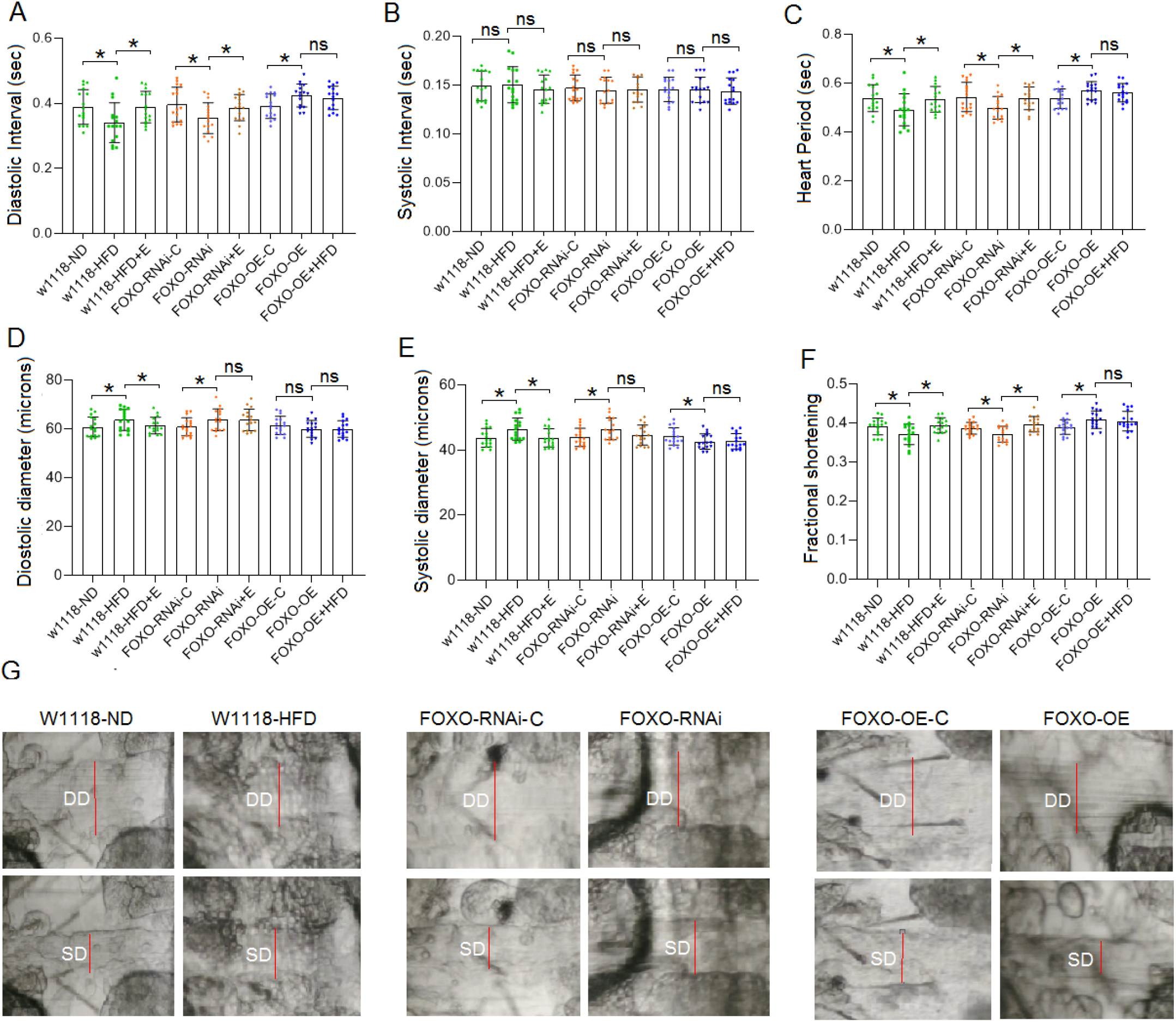
Heart function in 8-day old flies. (A) Heart diastolic period. (B) Heart systolic period. (C) Heart period. (D) Diastolic diameter. (E) Systolic diameter. (F) Fractional shortening. (J) Systolic image and diastolic image of flies. SD: systolic diameter; DD: diastolic diameter. For heart function, sample size was 17 hearts for each group, and the 1-way analysis of variance (ANOVA) with least significant difference (LSD) tests was used to identify differences among the groups with similar genetic backgrounds. Data are represented as means ± SEM. *P<0.05; **P <0.01.

The results showed that at 8 days of age, the diastolic diameter(DD) of w1118-HFD flies was significantly higher than that of w1118 flies(P<0.05), and the DD of w1118-HFD+E flies was significantly lower than that of w1118-HFD flies(P<0.05). Besides, the DD of FOXO-RNAi flies was significantly higher than that of FOXO-RNAi-C flies(P<0.05), but there was no significant difference in DD between FOXO-RNAi flies and FOXO-RNAi+E flies(P>0.05). Moreover, there was no significant difference in DD between FOXO-OE flies and FOXO-OE+HFD flies(P>0.05), and there was no significant difference in DD between FOXO-OE flies and FOXO-RNAi-C flies(P>0.05). (Fig.6-D and G)

The results showed that at 8 days of age, the systolic diameter(SD) of w1118-HFD flies was significantly higher than that of w1118 flies(P<0.05), and the SD of w1118-HFD+E flies was significantly lower than that of w1118-HFD flies(P<0.05). Besides, the SD of FOXO-RNAi flies was significantly higher than that of FOXO-RNAi-C flies(P<0.05), but there was no significant difference in SD between FOXO-RNAi flies and FOXO-RNAi+E flies(P>0.05). Moreover, the SD of FOXO-OE flies was significantly lower than that of FOXO-OE-C flies(P<0.01), and there was no significant difference in SD between FOXO-OE flies and FOXO-RNAi-UAS flies(P>0.05). (Fig.6-E and G)

The results showed that at 8 days of age, the fractional shortening(FS) of w1118-HFD flies was significantly lower than that of w1118 flies(P<0.05), and the FS of w1118-HFD+E flies was significantly higher than that of w1118-HFD flies(P<0.05). Besides, the FS of FOXO-RNAi flies was significantly lower than that of FOXO-RNAi-C flies(P<0.05), and FS of FOXO-RNAi+E flies was significantly higher than that of FOXO-RNAi flies(P<0.05). Moreover, the FS of FOXO-OE flies was significantly higher than that of FOXO-OE-C flies(P<0.05), but there was no significant difference in FS between FOXO-OE flies and FOXO-RNAi-UAS flies(P>0.05). (Fig.6-F)

The results showed that at 8 days of age, the heart TG level of *w*^*1118*^-HFD flies was significantly higher than that of *w*^*1118*^ flies(P<0.01), and the heart TG level of w1118-HFD+E flies was significantly lower than that of w1118-HFD flies(P<0.05). Besides, the heart TG level of FOXO-RNAi flies was significantly higher than that of FOXO-RNAi-C flies(P<0.05), and the heart TG level of FOXO-RNAi+E flies was significantly lower than that of FOXO-RNAi flies(P<0.01). Moreover, but there was no significant difference in heart TG level between FOXO-OE flies and FOXO-OE-C flies(P>0.05), and there was no significant difference in the heart TG level between FOXO-OE flies and FOXO-OE+HFD flies(P>0.05) (Fig.7-A and B).

**Fig.7.**
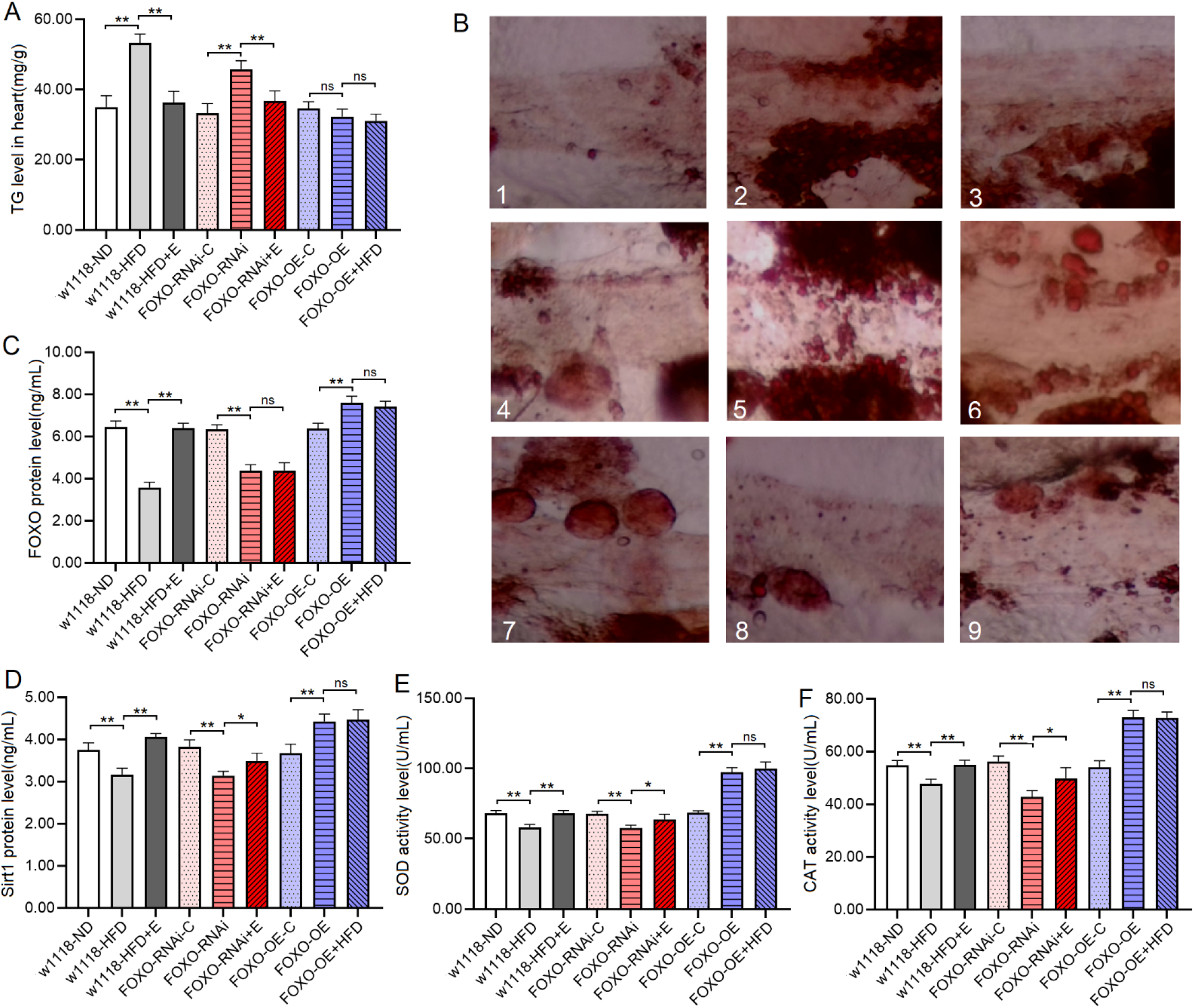
Heart TG level and heart Sirt1/FOXO pathways activity in 8-day old flies. (A) Heart TG level. (B) Oil red O staining of the heart of Drosophila. 1: w^1118^ flies; 2: w^1118^-HFD flies; 3: w^1118^-HFD+E flies; 4: FOXO-RNAi-UAS flies; 5: FOXO-RNAi flies; 6: FOXO-RNAi+E flies; 7: FOXO-OE-UAS flies; 8: FOXO-OE flies; 9: FOXO-OE+HFD flies. (C) Heart FOXO protein level. (D) Heart Sirt1 protein level. (E) Heart SOD activity level. (F) Heart CAT activity level. For protein measurement, the sample size was about 60 flies’ hearts for each group, and the 1-way analysis of variance (ANOVA) with least significant difference (LSD) tests was used to identify differences among the groups with similar genetic backgrounds. Measurements were taken 3 times. Data are represented as means ± SEM. *P<0.05; **P <0.01.

The proteins and activity of proteins showed that at 8 days of age, the heart FOXO protein level, Sirt1 protein level, SOD and CAT activity level of *w*^*1118*^-HFD flies were significantly lower than that of *w*^*1118*^ flies(P<0.01), and the heart FOXO protein level, Sirt1 protein level, SOD and CAT activity level of w1118-HFD+E flies were significantly higher than that of *w*^*1118*^-HFD flies(P<0.01). Besides, the heart FOXO protein level, Sirt1 protein level, SOD and CAT activity level of FOXO-RNAi flies were significantly lower than that of FOXO-RNAi-C flies(P<0.01), and the heart Sirt1 protein level, SOD and CAT activity level of FOXO-RNAi+E flies was significantly higher than that of FOXO-RNAi flies(P<0.01), but there was no significant difference in the heart FOXO protein level between FOXO-RNAi flies and FOXO-RNAi+E flies(P>0.05). Moreover, the heart FOXO protein level, Sirt1 protein level, SOD and CAT activity level of FOXO-OE flies were significantly higher than that of FOXO-OE-C flies(P<0.01), but there was no significant difference in the heart FOXO protein level, Sirt1 protein level, SOD and CAT activity level between FOXO-OE flies and FOXO-OE+HFD flies(P>0.05) (Fig.7-C to F).

The transmission electron microscope images displayed that a HFD and FSR disrupted the alignment of myofibrils and reduced the number of mitochondria, but EE protected them from damage induced by induced by HFD and FSR. FSO protected the alignment of myofibrils and the number of mitochondria from damage induced by HFD(Fig.8).

**Fig.8.**
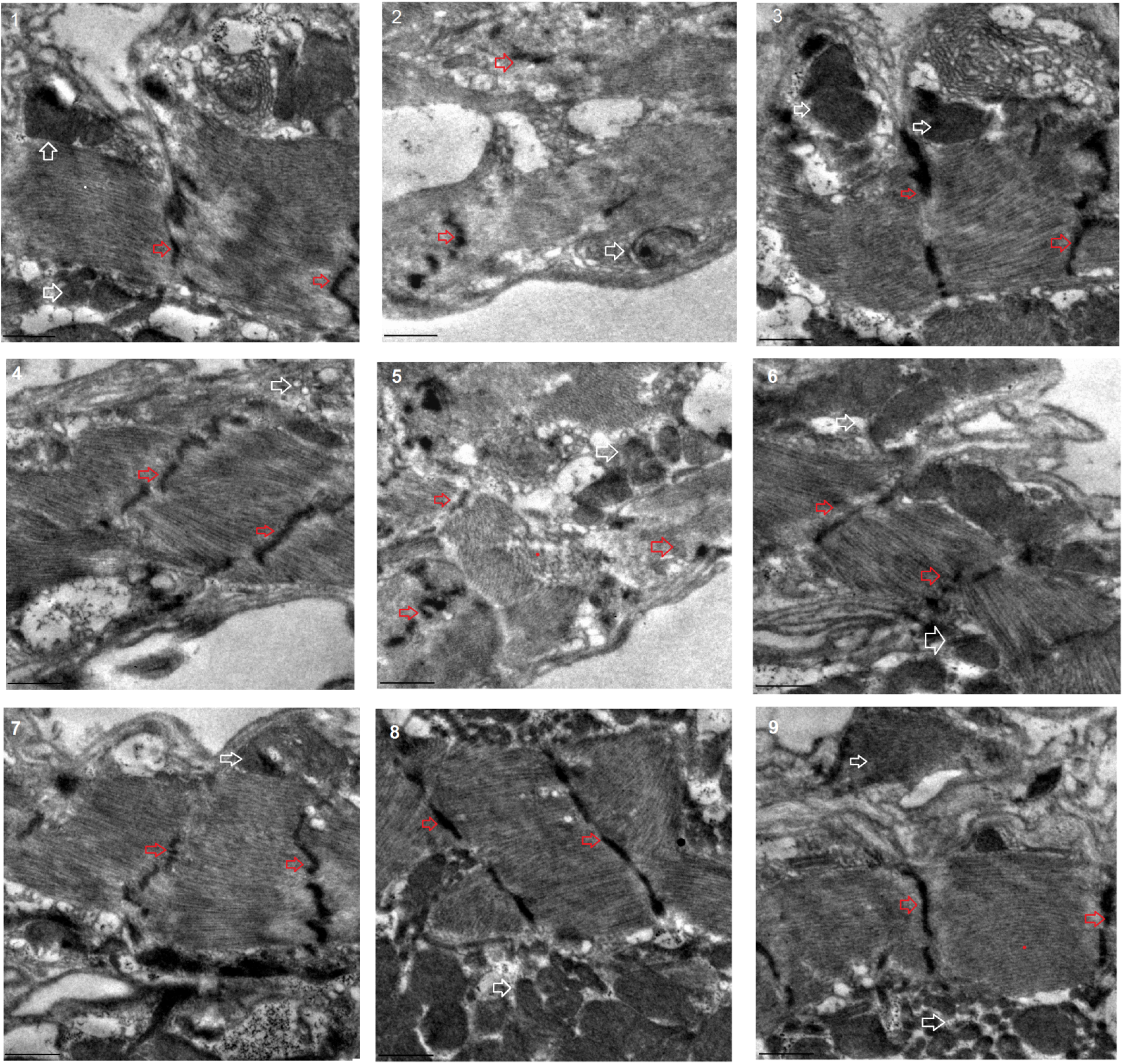
Transmission electron microscopy of heart. 1: w^1118^ flies; 2: w^1118^-HFD flies; 3: w^1118^-HFD+E flies; 4: FOXO-RNAi-UAS flies; 5: FOXO-RNAi flies; 6: FOXO-RNAi+E flies; 7: FOXO-OE-UAS flies; 8: FOXO-OE flies; 9: FOXO-OE+HFD flies. Scale: the black line represents 2 microns. The red arrow points to the Z line. The white arrow points to the mitochondria. The transmission electron microscope images displayed that HFD and FSR induced myofibrillary damage and mitochondrial damage, but PE could improve the myofibrillary damage and mitochondrial damage induced by HFD and FSR. FSO protected myofibrillary and mitochondrial from damage induced by HFD.

These results suggested that a HFD(with 15% lard) and MFSR induced cardiac defects, including weakening cardiac ability to contractility, increasing lipid accumulation and oxidative damage, inhibiting antioxidant capacity, and damaging myofibrils and mitochondria. However, EE and MFSR could prevent heart defects caused by a HFD(with 15% lard), and EE could prevent heart defects caused by MFSR. Therefore, cardiac FOXO gene played a key role in regulating HFD-induced cardiac dysfunction, which was consistent with the results of previous study[12], but FOXO was not a key gene for EE against heart defects caused a HFD in young flies, and its mechanism may be related to regulation of lipid catabolism and Sirt1/FOXO/SOD, CAT pathways related to antioxidant.

## 4 Discussion

*Drosophila* has emerged as an important model to study the effects of a HFD on metabolism, heart function, behavior, and ageing[39, 41–43]. For example, a HFD increases TG levels and disrupts insulin/glucose homeostasis in flies, which is similar to mammalian responses[44, 45], and it also causes cardiac TG accumulation, reduced heart contractility, conduction blocks, and severe structural pathologies, reminiscent of diabetic cardiomyopathies. Remarkably, these metabolic and cardiotoxic phenotypes induced by a HFD can be blocked by inhibiting insulin-TOR signaling[12, 17, 40]. Moreover, reducing insulin-TOR activity by expressing FOXO or *bmm* in myocardium alleviates cardiac TG accumulation and heart dysfunction induced by a HFD. Cardiac dysfunction induced by a HFD persists for two subsequent generations in *Drosophila*, and this is associated with reduced expression of *bmm* and *PGC-1α[12, 53, 54]*. Cardiac Nmnat/NAD^+^/SIR2 pathways are important antagonists of HFD-induced lipotoxic cardiomyopathy[55]. Endurance exercise(EE) could improve lipotoxic cardiomyopathy induced by a HFD or cardiac *dSir2* knockdown by activating NAD^+^/dSIR2/*PGC-1α* pathway in old *Drosophila*[17]. EE also improves exercise ability, cardiac contraction, and dSir2 expression, and it reduces TG levels, heart fibrillation, and mortality in both HFD and aging flies[37, 56]. These evidences suggest that FOXO pathways play a key role in the regulation of lipid metabolism in the heart of flies, but it remains unclear whether FOXO pathways mediate lipid metabolism regulation in skeletal muscle in young flies.

In this study, young male fruit flies were given a HFD intervention starting at 3 days of age for 6 consecutive days. The results showed that after feeding a HFD, the body TG level and the TG level of SM and HM were increased, and the bmm expression amd CPT1 activity were decreased, which indicated that lipid homeostasis had been disrupted. Besides, the results of climbing speed and climbing endurance suggested that a HFD impaired the exercise performance in young male flies, and the results of heart period and fractional shortening showed that a HFD impaired the heart function in young male flies. Further research found that a HFD reduced lipid catabolism and down-regulated Sirt1/FOXO/SOD, CAT pathways in skeletal muscle and heart of young male Drosophila, and it increased MDA level and damaged myofibrils in skeletal muscle and heart. These results indicated that the structure and function of SM and heart were impaired by a HFD, which may be related to the down-regulation of muscle FOXO/antioxidant pathways and lipid catabolic pathway.

In order to determine the regulatory function of *FOXO* gene on lipid metabolism in skeletal muscle and heart, *FOXO* gene was specifically overexpressed and knocked down in *Drosophila* SM and HM. The results showed that the structure and function of SM and heart were impaired by MFSR, and MFSR also down regulated muscle FOXO/antioxidant pathways and lipid catabolic pathway in skeletal muscle and heart of young male *Drosophila*, and it increased MDA level and damaged myofibrils in skeletal muscle and heart. These changes were similar to the skeletal muscle damages induced by a HFD. In contrast, muscle FOXO-specific overexpression(MFSO) not only reduced lipid accumulation in skeletal muscle and heart and improved exercise performance and heart function, but also effectively prevented HFD-induced defects in skeletal muscle and heart. MFSO up regulated FOXO/antioxidant pathways and lipid catabolic pathway in skeletal muscle and heart of young male Drosophila. Therefore, these results confirmed that FOXO played a vital role in regulating lipid metabolism in skeletal muscle and heart, and the mechanism is that FOXO in skeletal muscle and heart participated in the regulation of FOXO/antioxidant pathways and lipid catabolic pathway.

FOXO is a highly conserved transcription factor that plays an important role in cellular homeostasis. FOXO protein is also involved in the regulation of cell cycle, apoptosis and muscle regeneration[57, 58]. Silent information regulator 1 (Sirt1), a homolog of Sir2, appears to control the cellular response to stress by modulating FOXO, which acts as a sensor of insulin signaling pathways and a regulator of biological lifespan[59–61]. Sirt1 is an NAD^+^ dependent deacetylase and a commonly expressed protein that regulates gene expression through histone deacetylation, and it plays a complex role in the pathology, progression and treatment of a variety of diseases[62]. Sirt1 plays a key role in FOXO function through NADH-dependent deacetylation in oxidative stress response, which may contribute to stress resistance and longevity of cells[63]. In oxidative stress response, FOXO protein is translocated from the cytoplasm to the nucleus, promoting the expression of its target genes encoding antioxidant enzymes such as MnSOD, CAT, and glutathione peroxidase (GPX), and thus enhancing the antioxidant capacity of cells[64]. Therefore, Sirt1/FOXO/antioxidant pathways are involved in the regulation of lipid metabolism by regulating MDA and oxidative stress.

Excess lipids cannot be packaged into lipid droplets, resulting in a chronic rise in circulating fatty acids, which can reach toxic levels in non-adipose tissue. The deleterious effects of lipid accumulation in non-adipose tissue such as heart and skeletal muscle are known as lipotoxicity[5, 65–68]. Malondialdehyde (MDA) is the best product of lipid peroxidation, and it is produced from polyunsaturated fatty acids through chemical reactions and enzyme-catalyzed reactions. MDA is the most commonly measured biomarker of oxidative stress and lipid peroxidation[21, 69–71]. Moreover, The brummer (bmm) gene encodes the lipid storage droplet associated TG lipase Brummer, which is a homolog of human adipocyte triglyceride lipase (ATGL)[13, 72]. Lack of food or chronic overexpression of *bmm* depletes organismal fat stores in vivo, and loss of bmm activity causes obesity in flies[73]. Carnitine palmityl transferase I(CPT1) is the key enzyme in the carnitine dependent transport of long chain fatty acids across the mitochondrial inner membrane, and its deficiency results in a decrease rate of fatty acids beta-oxidation with decreased energy production[74–76]. FOXO gene overexpression limited the accumulation of lipids and increased *bmm* expression in heart[12]. Therefore, the regulation of *FOXO* on lipid metabolism is also related to the upstream regulation of *bmm*-related and CPT1-related lipid catabolic pathway.

Exercise is one of the best strategies to treat and prevent obesity[77–80]. There are many differences in long-term metabolic adaptations between endurance trained athletes and obese insulin-resistant volunteers, and examining the effects of chronic exercise training in diabetes-prone populations may reveal more interesting information about the role of lipotoxicity in the development of insulin resistance[65]. There is evidence that exercise can effectively counter lipotoxicity in skeletal and cardiac muscles[40, 67, 68]. On the one hand, exercise can increase fat consumption as energy, which can reduce fat accumulation and prevent obesity. Exercise, on the other hand, increases the antioxidant capacity of skeletal muscles and the heart, which reduces oxidative stress damage in skeletal muscle and heart[24, 81]. However, the role of FOXO gene in EE against lipid metabolism abnormalities and defects in heart and SM still remains unclear.

In this research, the results showed that in HFD flies, physical exercise decreased the body TG level, skeletal muscle TG level, and cardiac TG level, and it upregulated the bmm expression and CPT1 activity of skeletal muscle, which indicated that physical exercise protected lipid homeostasis from being disrupted by HFD. Moreover, the results of climbing speed and climbing endurance suggested that EE also protected exercise performance, heart period, and fractional shortening from being impaired induced by HFD in young male flies. Further research showed that physical exercise prevented the Sirt1/FOXO/SOD, CAT pathways from being down regulated by a HFD in young male, and it also prevented MDA level from being increased by a HFD, and EE protected myofibrils from being damaged by HFD in SM and heart. These results indicated that EE protected the structure and function from damages caused by a HFD in SM and heart, which may be related to the up-regulation of muscle lipid catabolic pathway.

Moreover, to determine whether FOXO played a key regulatory role in mediating EE resistance to defects induced by lipid accumulation, MFSR flies were also given an exercise intervention. The results showed that although EE did not upregulate FOXO gene expression and FOXO protein levels, it still upregulated lipid catabolic pathway and SOD, CAT activity, suggesting that FOXO was not a key regulatory gene of EE against lipid toxicity in SM and heart. Therefore, these results confirmed that EE prevented SM and heart from defects by up regulating Sirt1/antioxidant pathways and lipid catabolic pathway, but FOXO was not a key regulatory gene of EE against defects caused by a HFD in SM and heart.

In aged flies, cardiac *dSir2* gene knockdown induces lipotoxic cardiomyopathy via inhibiting cardiac NAD+/dSIR2/ *PGC-1α* pathway in old flies. A HFD aggravates lipotoxic cardiomyopathy induced by cardiac *dSir2* knockdown via inhibiting cardiac NAD+/dSIR2/*PGC-1α* pathway in untrained flies[55]. EE improve lipotoxic cardiomyopathy induced by cardiac *dSir2* knockdown, and it could prevent further deterioration of lipotoxic cardiomyopathy induced by a HFD in cardiac dSir2 knockdown flies[17]. Endurance exercise activated the cardiac Nmnat/NAD^+^/SIR2/FOXO pathway and the Nmnat/NAD+/SIR2/PGC-1α pathway, including up-regulating cardiac Nmnat, SIR2, FOXO and PGC-1α expression, superoxide dismutase (SOD) activity and NAD^+^ levels, and it prevented HFD-induced or cardiac Nmnat knockdown-induced cardiac lipid accumulation, malondialdehyde (MDA) content and fibrillation increase, and fractional shortening decrease[55].

## 5 Conclusion

Current findings confirmed that MFSO and EE protected SM and heart from defects caused by a HFD via enhancing FOXO-realated antioxidant pathways and lipid catabolism. FOXO played a vital role in regulating HFD-induced defects in SM and HM, but FOXO was not a key regulatory gene of EE against damages in SM and HM. The mechanism was related to activity of Sirt1/FOXO/ SOD,CAT pathways and lipid catabolism in SM and HM.

## Acknowledgements

We thank the Core Facility of Drosophila Resource and Technology, Center for Excellence in Molecular Cell Science, Chinese Academy of Sciences, for providing fly stocks and reagengts.

## Author Contributions

Research idea and study design: D.t.W.; data acquisition: D.t.W. X.K; data analysis/interpretation: D.t.W., Y.l.C; statistical analysis: D.t.W.; supervision: H.w.Q. Each author contributed during manuscript drafting or revision and approved the final version of the manuscript.

## Additional Information

Competing Interests: Authors have no conflicts of interest.

## Associated Data

### Data Availability Statement

All the generated data and the analysis developed in this study are included in this article.

